# The kisspeptin analog C6 reverses reproductive dysfunction in a mouse model of hyperprolactinaemia

**DOI:** 10.1101/2025.02.10.637587

**Authors:** Chloe Beaudou, Louise Sionneau, Didier Lomet, Vincent Robert, Peggy Jarrier Gaillard, Vincent Aucagne, Hugues Dardente, Massimiliano Beltramo, Vincent Hellier

**Affiliations:** INRAE, CNRS, Université de Tours, PRC, 37380, Nouzilly, France; Centre de Biophysique Moléculaire, CNRS UPR 4301, Rue Charles Sadron, 45071 Orléans, France

**Keywords:** Reproduction, Prolactin, Kisspeptin, Mouse

## Abstract

Hyperprolactinaemia (HPRL), characterized by elevated prolactin levels, disrupts the hypothalamic-pituitary-gonadal axis, leading to reproductive dysfunctions such as menstrual irregularities, anovulation, and infertility. Current treatments rely on dopamine agonists but are limited by side effects and resistance. Kisspeptin (Kp), a key neuropeptide regulating the reproductive function, offers potential as an alternative therapy. However, Kp’s short half-life requires impractical administration regimens. To address this, we developed a synthetic Kp analog, C6, with enhanced pharmacokinetics. This study evaluated the effects of C6 compared to Kp in a mouse model of HPRL. Mice received subcutaneous PRL injections for 21 days to induce HPRL, followed by daily or alternate-day intraperitoneal administration of Kp10, C6, or vehicle. Estrous cyclicity, luteinizing hormone (LH) secretion, ovarian histology, and hypothalamic gene expression were analyzed. As expected, the HPRL treatment blocked estrous activity, which was restored by both Kp10, the shortest bioactive isoform of Kp and C6. Histological analysis revealed increased corpora lutea in Kp10- and C6-treated groups, indicating restored ovulation. C6 demonstrated equivalent efficacy to Kp10 in mitigating HPRL-induced reproductive dysfunctions, offering a promising alternative therapy. Future investigations should further explore the mechanistic advantages of C6, particularly its role in LH regulation, to optimize treatment strategies for HPRL-related reproductive disorders.

## INTRODUCTION

Hyperprolactinaemia (HPRL), characterized by elevated levels of prolactin in the blood, presents a significant challenge to reproductive health, particularly in women. Prolactin (PRL), traditionally known for its role in lactation in a large variety of species^1^, exerts inhibitory effects on the hypothalamic-pituitary-gonadal (HPG) axis by suppressing the secretion of gonadotropin-releasing hormone (GnRH)^2,3^. In physiological conditions, lactation-induced elevated PRL secretion transiently inhibits the reproductive axis and prevents the occurrence of pregnancy. In pathophysiological conditions, HPRL in women leads to a long lasting inhibition of the reproductive system characterized by reduced GnRH and LH secretion, menstrual irregularities, anovulation, reduced sexual desire and infertility^4^. Given dopamine’s inhibitory effect on PRL release, medical treatments based on dopamine agonists, (e.g. bromocriptine or cabergoline)^5^ are currently the first line of treatment for patients suffering from HPRL. However, these drugs can induce pathopsychological side effects related to compulsive behaviours, and resistance^6^ can arise thus limiting treatment efficacy .

Acting upstream of GnRH neurons, neurons that express kisspeptin (Kp), a neuropeptide encoded by the KISS1 gene, have emerged as key regulators of the hypothalamo-pituitary-gonadal (HPG) axis^7^. Kp stimulates GnRH release ultimately driving gonadal function. Unlike GnRH, Kp neurons do express PRL receptors ^8–11^ and elevated levels of PRL in rodents are associated with a significant reduction of KISS1 mRNA expression ^11,12^. The working hypothesis is that HPRL effect on reproduction is caused by inhibition of Kp synthesis and release. This hypothesis is supported by both murine and human studies. Indeed, in murine models of HPRL, administration of the shortest active form of Kp composed by the ten last aminoacids, called Kp10, leads to normalization of estrous cyclicity, restoration of ovulation, and improved fertility^13^. Similarly, in clinical studies involving HPRL patients, intravenous perfusion or subcutaneous administration of Kp resulted in enhanced GnRH secretion, normalization of menstrual cycles, and improved ovulatory function^14,15^. However, due to the short half-life of the endogenous peptide^16^, a single dose of Kp is insufficient and either perfusion or multiple injections are required to achieve an effect. Such a dosing regimen is unsuitable for clinical applications. To palliate this issue we developed a synthetic analogue of Kp, called C6, retaining high affinity for the Kp receptor (KISS1R) and with enhanced pharmacokinetic properties compared to native Kp^17^. This analog has been shown to be active in several species ^17–21^ and could represent a potential alternative, or adjunctive therapy, for HPRL -associated reproductive dysfunctions. However, C6 effect on HPRL has not been tested yet. The aim of this study was to establish the possible advantage of C6 compared to Kp in rescuing reproductive function in a mouse model of HPRL. Treatments have been administered daily or every other day in a mouse model of HPRL and estrus cyclicity, LH plasma level, and ovarian histology were used as readouts for efficacy. Overall, we show here that HPRL induced reproductive dysfunction could potentially be reversed by C6 treatment and thus represent an interesting advance for possible therapeutic application.

## MATERIALS AND METHODS

### Mice and treatments

All procedures were performed according to EU legislation, were approved by the local ethical committee, and authorized by the French Ministry “*de l’enseignement supérieur et de la Recherche*” (authorization number: 2022101316093104). Nine-week-old C57Bl6/J mice were hosted 4 per cage with free access to food and water under a 12 hours light/dark cycle. After a period of habituation, they received twice daily a subcutaneous (s.c.) injection of prolactin (NIDDK-oPRL-19, AFP-9221A) at a dose of 7µg/day (dissolved in 18% ethanol 100% in phosphate buffer 0.1M; HPRL group) or vehicle (18%Eth/PB) (Control group) during 21 days. Control and HPRL mice groups were further subdivided in three different groups receiving daily intraperitoneal (i.p.) injections of either mouse Kp10 (1nmol, GeneCust), C6 (1nmol, synthesized in the lab) or DMSO (1% in saline solution, D4540 Merck), the vehicle of Kp10 and C6 solutions. Kp10- and C6-treated HPRL groups were subdivided in two additional groups receiving every two days an i.p. injection of either Kp10 (1nmol) or C6 (1nmol). Kp, C6 or DMSO treatments started on the same day as PRL injections.

### Analysis of estrous cyclicity

The estrous cycle stage of female mice was determined by vaginal smear following the method described by McLean AC et al, 2012^22^. Smears were taken each day during 20 days, approximately 2h after the beginning of the light period. Analysis of the presence of either lymphocytes, nucleated epithelial cells or keratinocytes was performed in a blinded manner by the same experimenter that assigned the status of the mouse to one of the different oestrous phases: diestrus, metestrus, proestrus, and estrus.

### LH ELISA

After the 21 days period of the protocol, at 9:00 am, Kp10 and C6 were administered to the daily treated groups, others groups received a saline injection. Mice receiving the treatment every two days were injected with Kp10 or C6 the day before. Blood samples (4μL) were collected every 20 minutes during 2 hours starting at 9:00am after an initial and unique section of 1mm of the tail^23^. A sandwich ELISA, adapted from Steyn et al.^23^ and previously described ^17^, was used to measure LH concentration in blood. Monoclonal bovine LHβ 518 B7 (1:1,000 dilution) was used as the capture antibody (obtained from Lillian Sibley at University of California Davis), and the standard was NIDDK mouse LH reference preparation from AF Parlow (AFP5306A mouse RIA kit). The assay sensitivity was 0.2 ng/mL and the mean intra-assay coefficient of variation averaged 11%.

### Prolactin ELISA

Twenty-two days after the beginning of the protocol, blood samples were collected after decapitation and PRL measured by ELISA method (E0022Sh, BT Lab) following manufacturer’s instructions. The assay sensitivity was 2.54ng/mL. Mice receiving 7µg/day injections of PRL induced a PRL serum concentration of 225.6 ± 9.9ng/mL.

### Ovaries treatment

Twenty-two days after the beginning of PRL treatment, mice were decapitated and ovaries were sampled. Samples were immersed in PBS and fixed for 4 hours in 4% PFA, washed, paraffin-embedded and sectioned at 4μm. Sections were stained with hematoxylin and eosin Y. All sections were collected and images acquired with an AxioScanZ1 slide scanner (Carl Zeiss, France). Both ovaries of each animal were examined and the number of corpora lutea counted in all ovaries.

### Gene expression analysis

Brains were collected at the time of sacrifice (see “Ovaries treatment section”). RNA extraction was realized from hypothalamic blocks containing the median eminence, arcuate nucleus, dorsomedial/ventromedial hypothalamic nuclei and RT-qPCR were performed as described previously (see Abot et al 2022^24^). RNA was extracted using TriReagent (Sigma). Concentration and purity of individual samples were determined with Nanodrop 2000 (ThermoScientific) and integrity was checked by standard agarose gel electrophoresis. cDNA was synthesized using Omniscript RT kit (Qiagen) and Oligo-dT primers (Eurofins, Germany). As a negative control, the same mix with water instead of RT was prepared. Quantitative PCR was performed using CFX-96 Real-Time PCR Detection System (BioRad) and Sso Adv Universal SYBR Green Supermix (Bio-Rad). All samples (unknown, standard curves) were assessed in triplicate and Rplp0 (a.k.a 36B4) was used as a housekeeping gene to normalize expression. All primer pairs (table 1) had an efficiency of ∼90-110% and a single sharp melting peak was observed. The quantification of mRNA level was obtained by the 2-ΔΔCT method and data are presented as fold-increase compared to the control condition.

**Table 1.**
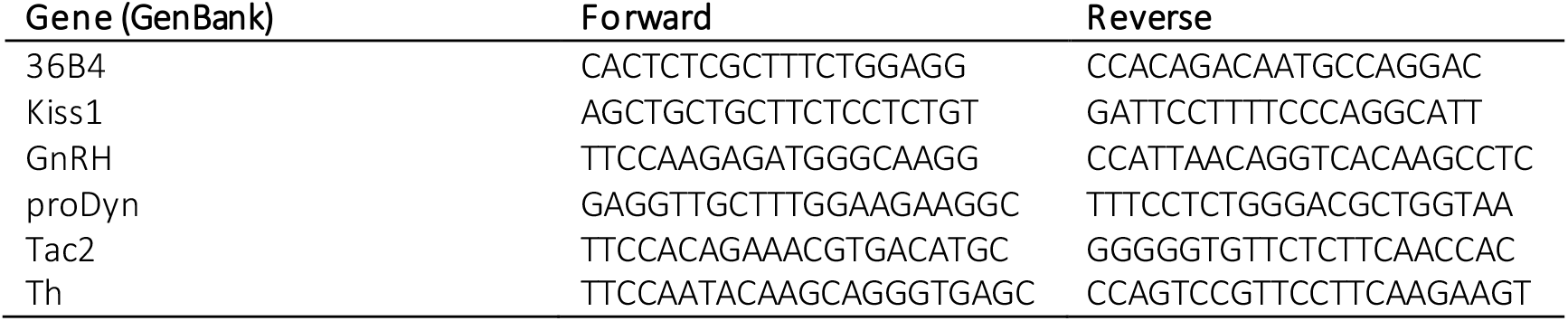
List of primers used for qRT-PCR.

### Statistics

Data were analysed using GraphPad Prism 10.2.3 and are expressed as box and whiskers. Differences between groups were analysed using the Kruskal-Wallis statistical test followed by Dunn’s post hoc test. P values lower than 0.05 were considered statistically significant.

## RESULTS

To address the effects of Kp or C6 treatment on HPRL syndrome, we developed a HPRL mouse model by injecting PRL subcutaneously twice daily during 21 days (Fig. 1A). In mice, an estrus cycle is characterized by the succession of 4 phases : metestrus (24h), proestrus (less than 24h), estrus (less than 24h) and diestrus (approximately 2 days)^25^. As expected, control mice showed 4 to 5 days regular estrus cycles while HPRL mice, after the 7^th^ day of treatment and until the end of the injection period (21 days), exhibited abnormal or absent estrus cyclicity (Fig. 1B). However, daily or every two days injection of Kp10 or C6 induced an improvement of the regularity of the estrus cyclicity (Fig. 1C). Overall, HPRL mice spent significantly more time in metestrus compared to control mice (p<0.0001) (Fig. 1D), and exhibited less estrus phase (Fig. 1E). Interestingly, the percentage of time spent in metestrus by HPRL mice upon C6 treatment (either injected daily or every two days) is not significantly different from control mice (p=0.2792 and p=0.2560 respectively). Conversely, Kp-10-treated mice spent significantly more time in metestrus compared to control DMSO-treated mice (daily injection: p=0.008; every two days injection p=0.01)(Fig 1D). However, the percentage of time spent in metestrus between HPRL group and HPRL-Kp10- or C6-treated groups (either injected daily or every two days) was not statistically different. Both Kp10 and C6 treated mice showed an improvement of estrus cyclicity with an increase in the percentage of mice showing at least one estrus, compared to HPRL mice treated with vehicle (Fig. 1E).

**Figure 1.**
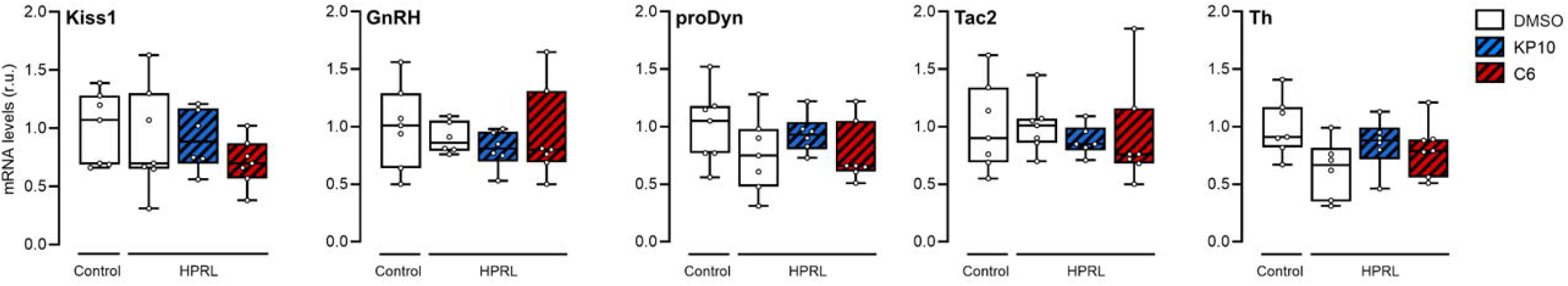
HPRL mouse model and effect of treatments on estrous cyclicity. (A) Mice are receiving twice a day an injection of prolactin at a dose of 7ug/day (HPRL group) versus vehicle (Control group). An additional treatment is administered daily (Kp10, C6 or DMSO group) or every two days (Kp10 -2D or C6-2D group) in order to evaluate the effect of Kp10 and C6 treatment. (B-C) Representative graphs of estrous cyclicity in the Control and HPRL groups (B) and upon treatments with Kp10 or C6 (C). (D) Percentage of time spent in the metestrus phase of the estrus cycle. (E) Percentage of mice showing at least one estrus after the 7^th^ day of treatment. Data are represented as box and whiskers. n=between 9 and 11 per group; * p<0.05; **p<0.01; ****p<0.0001 (Kruskal-Wallis test followed by a Dunn’s post-hoc analysis).

At the end of the 21-day treatment period, Kp10 or C6 have been injected to evaluate the effect of chronic treatments on LH secretion after bolus injection (Fig. 2A). Of note, daily treated mice received an injection of respective treatments on the day of blood sampling. Every two days treated mice were injected the day before the blood sampling. Blood sampling was performed every 20 minutes during 2 hours. Daily injections of both treatments did not affect LH secretion in control mice (Fig. 2B). However, C6 (but not Kp10) injection every two days had a significant effect on LH secretion of HPRL mice in comparison to control HPRL mice (p=0.0087) (Fig. 2C).

**Figure 2.**
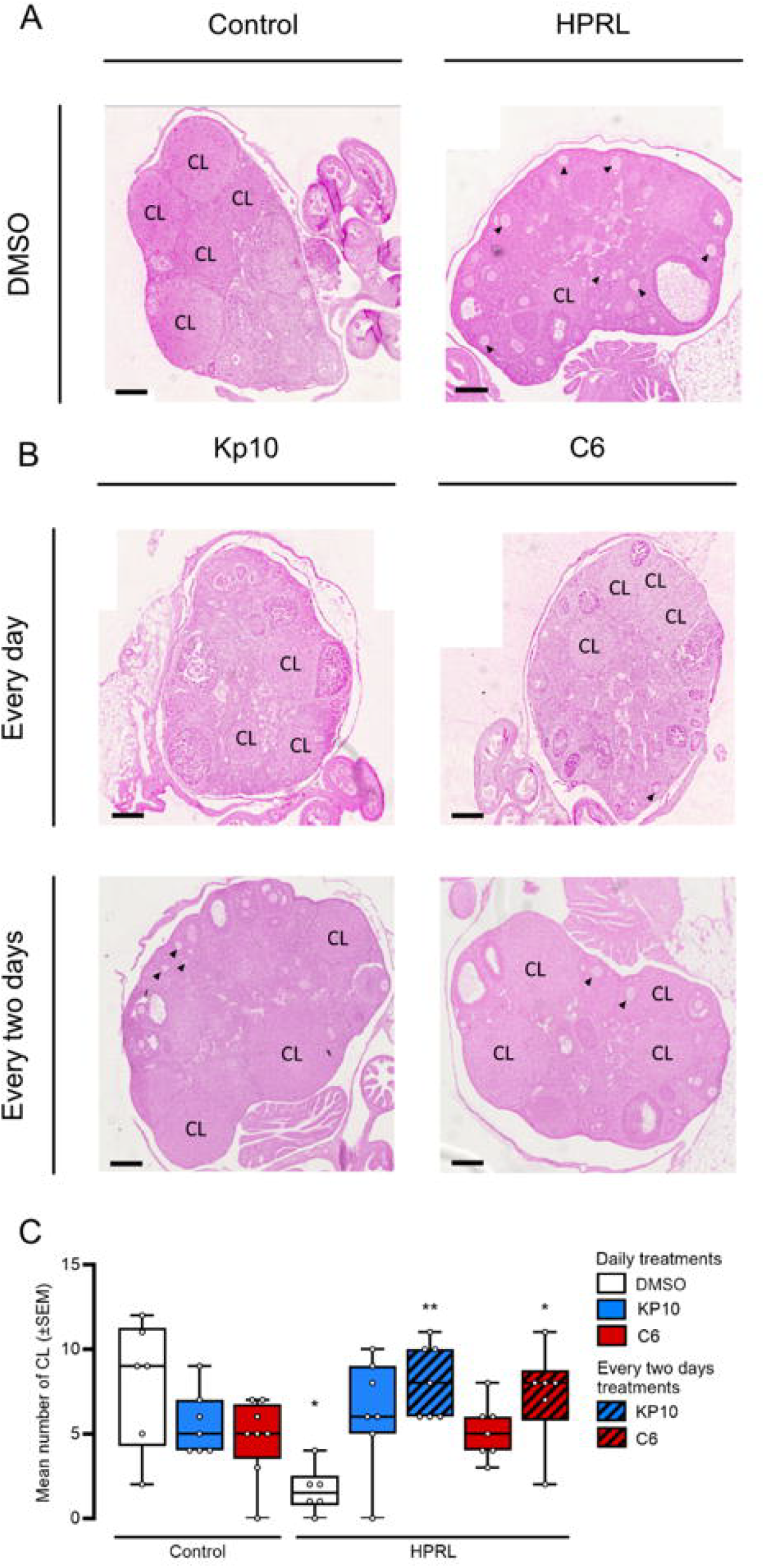
Effect of Kp10 and C6 treatments on LH secretion. (A) Schematic representation of the 21 days (one dotted line representing one day) of treatment, with an administration every days or every two days (black dots) and serial blood sampling over 120 minutes in HPRL and control mice at the end of the 21 days upon administration of the different treatments. (B-C) Evaluation of the total amount of LH secreted (AUC) upon DMSO, Kp or C6 administration everyday (B) or every two days (C). Results are represented as AUC as box and whiskers. n=between 9 and 11 per group. **p<0.01 (Kruskal-Wallis test followed by a Dunn’s post-hoc test)

Gene expression analysis on medio-basal hypothalamus (MBH) from HPRL and control mice upon 21-day of treatment revealed no difference between groups regardless of treatment (Fig. 3). To gain further insight into the effect of Kp10 and C6 treatment on the reproductive function of HPRL mice, ovaries from HPRL and control mice were collected and analysed. As shown in figure 4, HPRL mice showed a significant reduction of the number of corpora lutea as compared to control mice (p=0.0180; Kruskal-Wallis). Of note, histological observation of the ovaries of HPRL mice revealed the presence of numerous primary follicles (Fig. 4A-B). In HPRL mice, Kp10 or C6 treatments provided every other day, but not daily, increased the number of corpora lutea compared to control HPRL mice (p=0.0044 and p=0.0326 respectively) (Fig. 4B-C).

**Figure 3.**
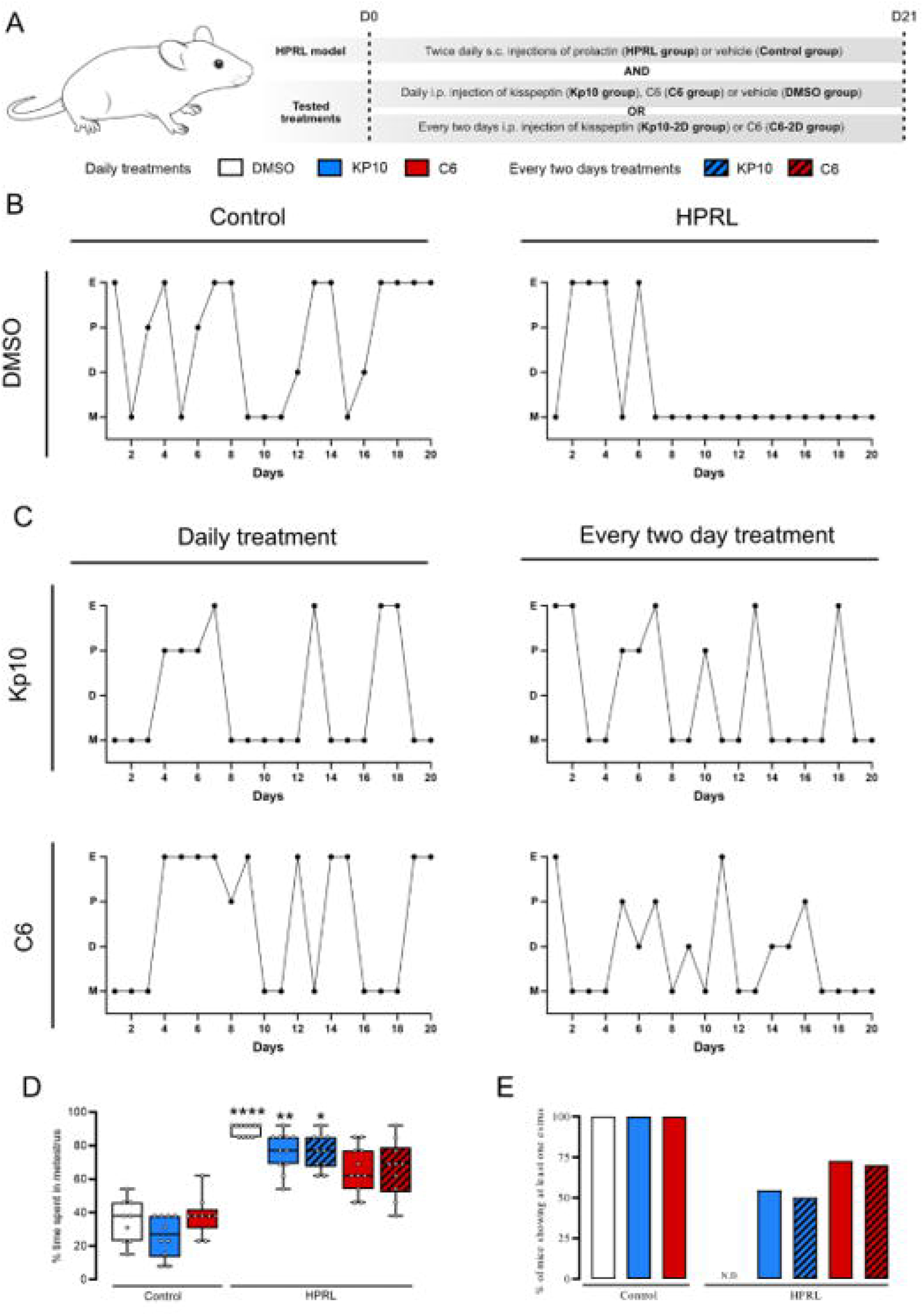
Expression of Kiss1, GnRH, proDyn, Tac2 and Th genes in the medio-basal hypothalamus of HPRL mice upon KP10 or C6 treatments every two days.

**Figure 4.**
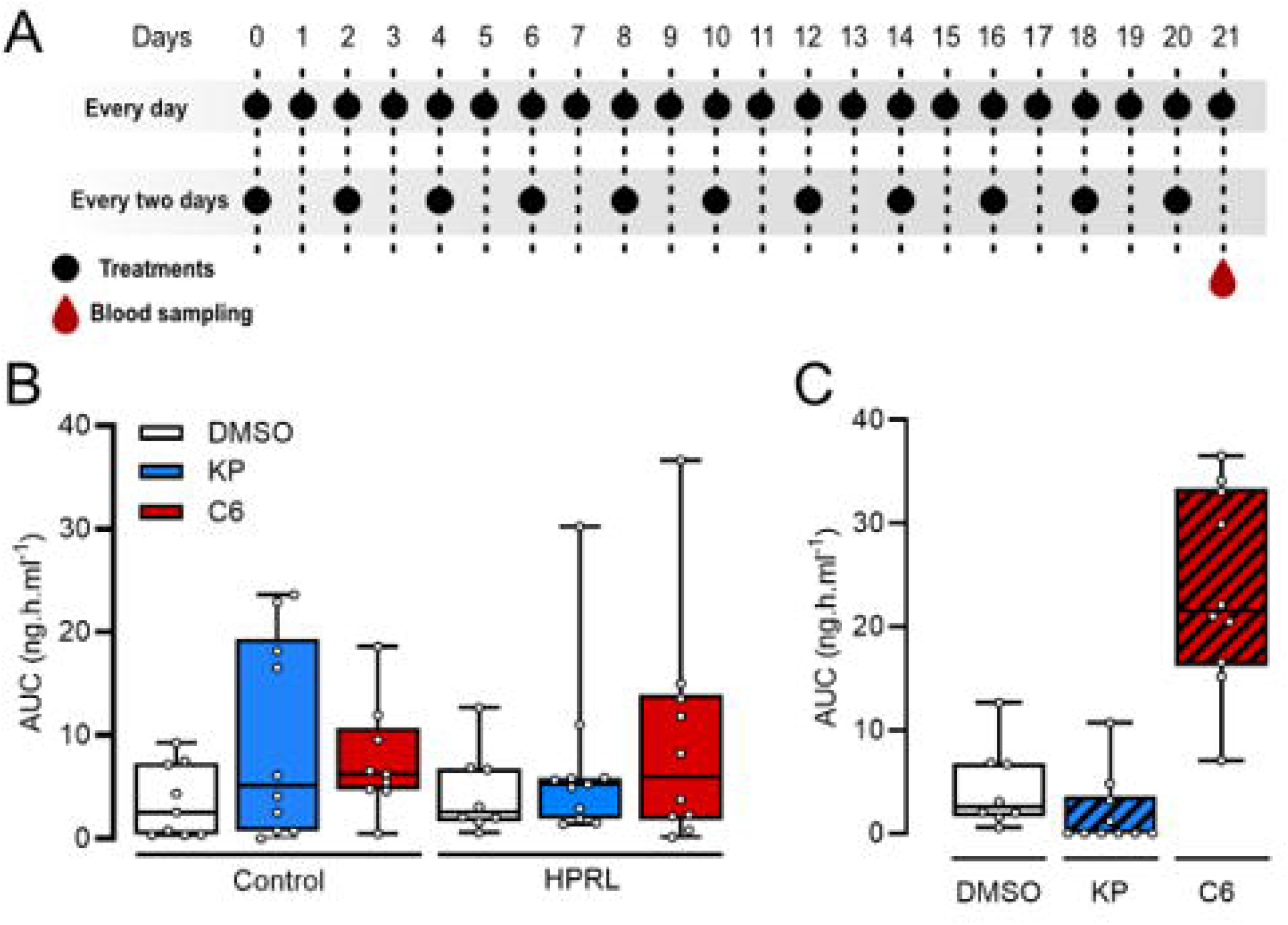
Ovarian histology (A-B) and number of corpora lutea (C) upon Kp10 and C6 treatment on HPRL mice. Arrows in A and B indicate primary follicles. Scale bar 150μm. CL: Corpus luteum. n=between 6 and 8 per group *p<0.05; **p<0.01 Kruskal-Wallis followed by a Dunn’s post-hoc test.

## DISCUSSION

Here we show that C6, a Kp analog developed in our laboratory, reduced the negative reproductive impact of PRL in a mouse model of HPRL. As previously reported^13^, Kp10 treatment restored estrus cyclicity over a 21 days period. C6 treatment elicited a similar effect. Under a protocol of PRL daily administration, both compounds, regardless of administration frequency, improved estrous cyclicity (increasing the time spent in metestrus) and estrus occurrence. As biomarker of reproductive physiology status, we analysed LH secretion in the different groups of mice. Control and HPRL mice exhibited low LH levels which is consistent with a lack of stimulation on one hand and inhibition induced by PRL administrations on the other hand. At the end of the daily treatment period, neither Kp nor C6 injection elicited a significant increase of LH blood level. Clinical trials in women suffering from reproductive dysfunction and treated twice daily with Kp showed that such a pattern of administration led to receptor desensitization and an interruption of the stimulatory effect of Kp on gonadotropin secretion^26^. Additional experiments led in sheep revealed that continuous administration of Kp prevented the effect of the peptide on LH secretion over time, which is in accordance with a desensitization of the receptor^27^. Finally, studies in male rats revealed similar LH releasing profiles upon continuous subcutaneous administration, with a reduction of LH secretion within 24 hours^28,29^. Taken together, these observations suggest that daily administration of Kp or C6 in our protocol could potentially lead to receptor desensitization and an absence of stimulation of LH release.

More interestingly, every two days administration of Kp or C6 led to divergent effects on LH release. C6-treated mice showed an elevated LH level, whereas in Kp-treated mice LH levels were similar to that of the control group. The different half-life of Kp10 and C6 could help explain this result. Indeed, with a very short half-life ^16^, every two days injection of Kp may not be sufficient to have a long lasting effect on LH release. However, C6, with a half-life of several hours, seems to induce a marked increase of LH levels in the plasma. This may be due to the capacity to reach a steady state concentration of C6 that is not inducing receptor desensitisation. However, continuous LH release is not ideal to induce physiological stimulation of ovulation. Hence, a different dosing regimen like a 3 or 4 days injection may be more appropriate to prevent this supraphysiological stimulation of LH release observed in our study. Another possibility may be that C6, but not Kp10, would induce some transcriptional events leading to a long-term change in gene expression. However, expression level of genes known to be involved in the control of LH secretion, and more generally, in the control of reproduction does not seem to be influenced by these treatments. Injected peripherally, Kp10 and C6 peptides are unable to reach the brain^30^ (and unpublished data). Hence, their effect at the central level appears to be restricted to brains regions devoid of the blood brain barrier. This would prevent direct effects on neurons expressing the key genes analysed in this paper.

To characterize further the impact of the treatments, we performed histological analysis of ovaries. Overall, treatments with Kp and C6 induced a significant increase in the number of corpora lutea compared to HPRL mice control group. In contrast to the effect on LH release, daily and every two days treatments with either molecule elicited the same effect on ovulation and mirrored the effects observed on estrus cyclicity. These results suggest a potential dichotomy between central versus peripheral effect of the treatments. In line with this hypothesis, recent findings revealed a potential direct effect of Kp on the ovaries. Indeed, expression of the Kiss1 receptor has been detected in the ovary of several species^31–33^ and oocyte-selective ablation of Kiss1R leads to ovarian failure without significant effect on puberty onset^32,34^. Although these studies have not been performed with C6 yet, an ovarian effect of C6 cannot be excluded. Hence, our results may be explained by a distinct direct action of Kp and C6 in the ovaries which could lead to a stimulation of ovulation.

Due to the central role of Kp in reproduction, pharmacological regulation of the female HPG using Kp or its analog C6 is of high interest. Interestingly, the endogenous Kp10 peptide acts upon different parameters of the reproductive physiology, from ovulation to sexual desire and behaviour^35,36^, which presents substantial therapeutic potential in a wide range of reproductive disorders. To overcome biological limitation of short-lived Kp10 activity, we developed C6, an analog with similar effect on puberty, ovulation and LH release in various species^17–21^, but with improved half-life. We show that, similar to Kp10, C6 produces a reversal of HPRL ovarian cycle disruption. In addition, depending on the dosing regimen, it has an additional effect on LH secretion. The potential advantage of this effect on LH should be further investigated to fully understand its usefulness in the treatment of HPRL.

## Data availability

Data will be made available on request.

## Author contribution

**Vincent Hellier:** Writing – review & editing, Writing – original draft, Project administration, Investigation, Funding acquisition, Formal analysis, Conceptualization. **Chloe Beaudou, Louise Sionneau:** Writing, investigation and data curation. **Didier Lomet, Vincent Robert, Peggy Jarrier Gaillard:** Investigation. **Vincent Aucagne, Hugues Dardente:** Writing – review & editing. **Massimiliano Beltramo :** Funding acquisition.

